# Temperature Dependence of the *Krokinobacter* rhodopsin 2 Kinetics

**DOI:** 10.1101/2020.07.24.219782

**Authors:** P. Eberhardt, C. Slavov, J. Sörmann, C. Bamann, M. Braun, J. Wachtveitl

## Abstract

Here we applied target analysis to a temperature dependent flash photolysis dataset of the light-driven sodium ion pump *Krokinobacter* rhodopsin 2 (KR2) at sodium pumping conditions. With an increase in temperature from 10 – 40 °C, the overall photocycle duration was accelerated by a factor of six, while single transitions like the L to M transition increased by a factor of 40. Using kinetic modeling with the Eyring constraint as well as spectral corrections on the datasets the spectral position as well as the equilibria of the different photointermediates could be resolved. The results provide further insight into KR2s photocycle and energetics.

**STATEMENT OF SIGNIFICANCE:** KR2 is the most prominent member of the new class of non-proton cation pumps, as it represents an interesting new optogenetic tool. Despite widespread biophysical investigations, the molecular mechanisms of light-induced sodium pumping in KR2 are still not sufficiently understood. Therefore, an expanded set of thermodynamic parameters is essential for a complete picture. Our study of the KR2 photocycle shows that different steps in the photocycle are affected differently by temperature changes. Rigorous data analysis provides strong evidence that the transient states observed in time-resolved experiments represent rather equilibria between the different photocycle intermediates than pure intermediates. Gaining access to the dynamics and energetics of KR2 helps to answer long standing open questions concerning the molecular mechanism of cation pumping.

## INTRODUCTION

Rhodopsins are currently considered to be the most abundant light harvesting proteins on earth^1^. They have already been identified in all three domains of life (archaea, eubacteria and eukarya)^2–4^ as well as giant viruses^3,5^. Microbial rhodopsins generally consist of a seven transmembrane α-helix motif (opsin) and a covalently bound retinal chromophore. They fulfill a wide variety of functions ranging from light-energy conversion to signaling. Pump- and channel-type rhodopsins have gained significant attention due to their applications in optogenetics.^6–8^

In 2013 Inoue *et al.* described a new microbial rhodopsin from *Krokinobacter eikastus*, the *Krokinobacter* rhodopsin 2 (KR2) which is the first light driven outward Na^+^ pump discovered.^9^ KR2 can also pump H^+^ and Li^+^ in the absence of Na^+^ ions. Since Na^+^ ions can be used for neuronal silencing, Na^+^ pumps are of great interest for optogenetic experiments.^7,10^ Photon absorption in KR2 triggers *all-trans* to *13-cis* isomerization of the bound retinal.^11,12^ Subsequently, the protein undergoes a cyclic transformation through a sequence of different photointermediates, which is known as the photocycle. Usually, every photointermediate exhibits a unique spectral fingerprint, which can be used to describe and model the photocycle. Extensive studies have been performed on the first known microbial rhodopsin, bacteriorhodopsin (BR)^13^ to disentangle its rather complex photocycle. While at first, it was modeled as an unbranched unidirectional photocycle with five distinct photointermediates,^14^ later on more complex models were proposed.^15,16,17^ KR2’s photocycle has already been studied extensively with various techniques^18–23^ but still remains to be understood completely. Especially the spectral location of the photointermediates, as well as the equilibria within the photocycle remain elusive.

## MATERIALS AND METHODS

### Sample preparation

The KR2 gene (pet21 plasmid) was heterologously expressed in *Escherichia coli* cells C43(DE3) in mineral medium M9. Protein expression was induced with 0.5 mM isopropyl β-D-thiogalactopyranoside (IPTG; Carl Roth GmbH, Karlsruhe, Germany) and 5 µM all-trans retinal (ATR; Sigma-Aldrich, St. Louis, USA) was added. The protein was isolated by affinity chromatography (5 ml HisTrap, GE Healthcare Life Science, Chicago, USA) and desalted (5 ml HiTrap, GE Healthcare Life Science, Chicago, USA). KR2 was incorporated into liposomes with an inside-out orientation. The liposomes contained POPC, POPG and cholesterol in a ratio 8:1:1 and the lipid-protein-ratio was ∼6.

### Temperature dependent broadband flash photolysis measurement

Measurements were performed with a home-built setup as described previously.^24^ The sample was resuspended in buffer containing 50 mM NaCl and 50 mM TRIS at pH 8.5. The solution was placed in a 2 x 10 mm quartz cuvette in a tempered cuvette holder. The temperature of the cuvette holder was maintained by a thermostat (Pilot ONE, Huber, Offenburg, Germany). The sample was excited by a Nd:YAG laser (SpitLight 600, InnoLas Laser, Krailling, Germany) pumping an optical parametric oscillator (preciScan, GWU-Lasertechnik, Erfstadt, Germany). The optical parametric oscillator was set to generate single nanosecond pulses with a central wavelength of λ_max_ = 525 nm. White probe light was generated by a spectrally broad xenon flash lamp (model No. L7685, Hamamatsu, Hamamatsu City, Japan) and detected by a fast, intensified charge-coupled device camera (PI-MAX 3, Princeton Instruments, Beijing, China). The probe light was split before passing the sample to also record a reference beam. Spectra were collected between 1 µs and 500 ms.

### Data Analysis (Target Analysis)

The analysis of the flash photolysis datasets was performed using OPTIMUS (www.optimusfit.org).^25^ First, the datasets were subjected to lifetime distribution analysis, which is a model independent method. In this analysis, the pre-exponential amplitudes of a set of 100 exponential functions with fixed, equally spaced (on a decimal logarithm scale) lifetimes are determined. The obtained pre-exponential amplitudes at each detection wavelength can be presented in the form of a contour lifetime density map (LDM). The reading of the LDMs is as for a decay-associated spectrum from global lifetime analysis: (i) positive amplitudes account for decay of absorption or rise of ground state bleach (GSB); (ii) negative amplitudes account for rise of absorption or decay of GSB.

In addition, global target modeling was performed as described previously.^25^ All datasets recorded at different temperatures were fitted simultaneously within the same fitting routine and sharing one set of kinetic rates. The rates at one temperature were optimized non-linearly, while the rates at the other temperatures were calculated based on the Eyring equation (eq. 1, 2).

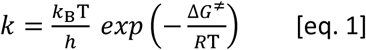

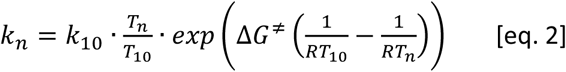

With this approach we fully employ the potential of global analysis^25,26^ and reduce the number of non-linear fit parameters for all datasets to one set of kinetics rates and the free enthalpy of activation ΔG^≠^. Moreover, we impose a restriction on the viable models that they should follow the Eyring theory, thereby eliminating a great of number of mathematically feasible but physically unreasonable models. Further advantage of the target analysis is that it also yields the spectra of the model states, which are helpful for further analysis and interpretation of the data.

### Spectral decomposition

The spectra obtained from the target analysis at a given temperature were fitted simultaneously with a set of four Gaussian functions with the width of the Gaussian function and its central wavelength as fit parameters. This procedure was performed independently for every temperature to also account for possible temperature effects on the spectra. Regardless, the fitted values showed only a minor temperature variation (∼ 0.6 % for the spectral position and ∼ 7 % for the width). The analysis was performed on the energy scale.

## RESULTS & DISCUSSION

### µs-ms kinetics of KR2

As an example of the KR2 photocycle kinetics a flash photolysis data set from a room temperature (20°C) measurement is shown in Figure 1 on the left. The positive signal present at the start of the measurement in the region between 550 – 650 nm is attributed to the K intermediate of KR2. At the same time, until the end of the photocycle, the negative signal of the ground state bleach (GSB) can be observed in the region between 460 and 550 nm. Following the K intermediate, the L intermediate is superimposed on the spectral region of the bleach, roughly centered around 500 nm. With the decay of the L intermediate, the blue shifted M intermediate is formed. Its very weak signal can be found between 400 and 475 nm around 0.1 – 1 ms. The last photointermediate is the red shifted O intermediate which is formed with the decay of M and occupies the spectral range between 550 and 670 nm. The O intermediate as well as the GSB decay with the lifetime of 6 ms, marking the end of the photocycle. Since the GSB and all the photointermediates have different spectral overlaps, which can be observed by the shifting behavior of the GSB, it is difficult the exact spectral characteristics of the photointermediates. Figure 1 on the right shows the same photocycle at 40 °C. Here, comparing it to the 20 °C photocycle one can observe the acceleration of the photocycle at higher temperatures. Especially prominent is the acceleration of the K intermediate, which is barely visible in the 40 °C experiment because it decays much faster than at 20 °C.

**Figure 1:**
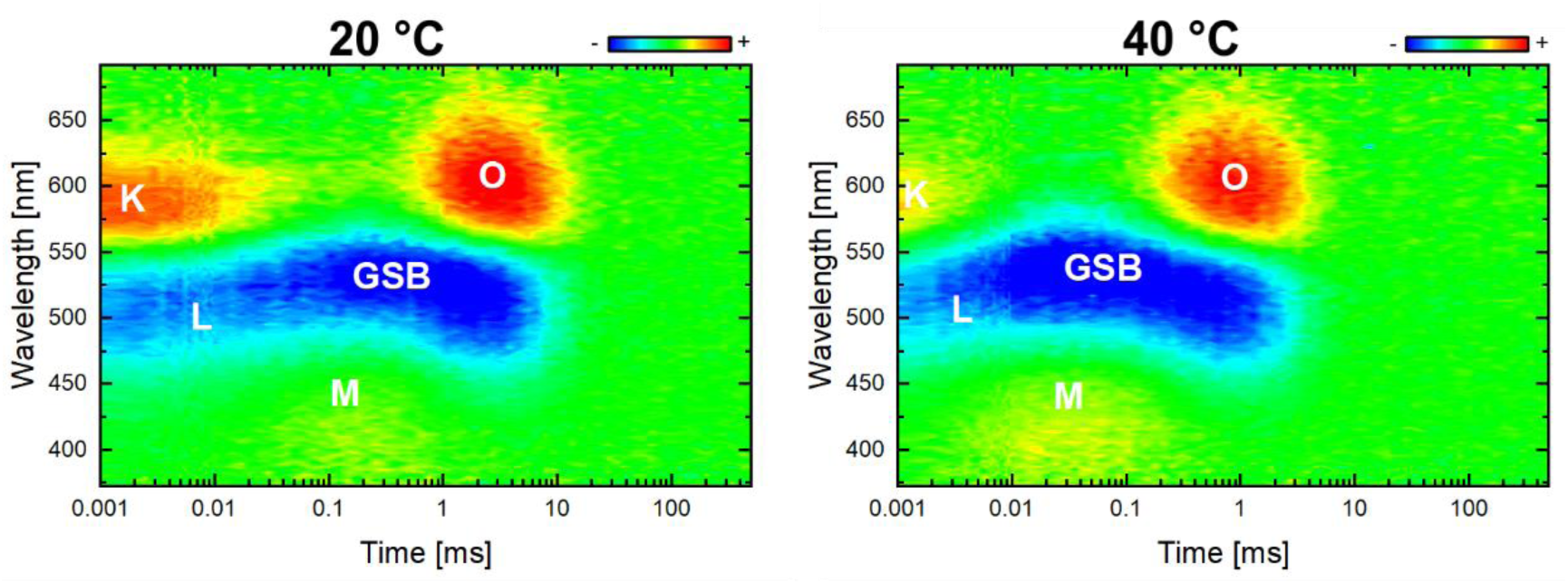
Broadband flash photolysis measurement of the KR2 wildtype with 50 mM NaCl, pH 8.5 at 20 °C (left) and 40 °C (right). The amplitudes are color coded. Red, green and blue indicate positive, zero and negative absorbance changes, respectively.

### Temperature dependence of the KR2 photocycle

To gain further insight in to the KR2 photocycle and the spectral position of the photointermediates we recorded time-resolved, temperature dependent flash photolysis data. Figure 2 shows the time traces of the absorbance changes of KR2 wildtype at four distinct wavelengths, 410, 460, 525 and 610 nm, as well as seven different temperatures (10 to 40 °C in steps of 5 °C). No excitation wavelength dependence (excitation at 480, 525 and 570 nm) of the flash photolysis data was observed (Figure S1), and thus inhomogeneity of the kinetics was not further considered in this study.

**Figure 2:**
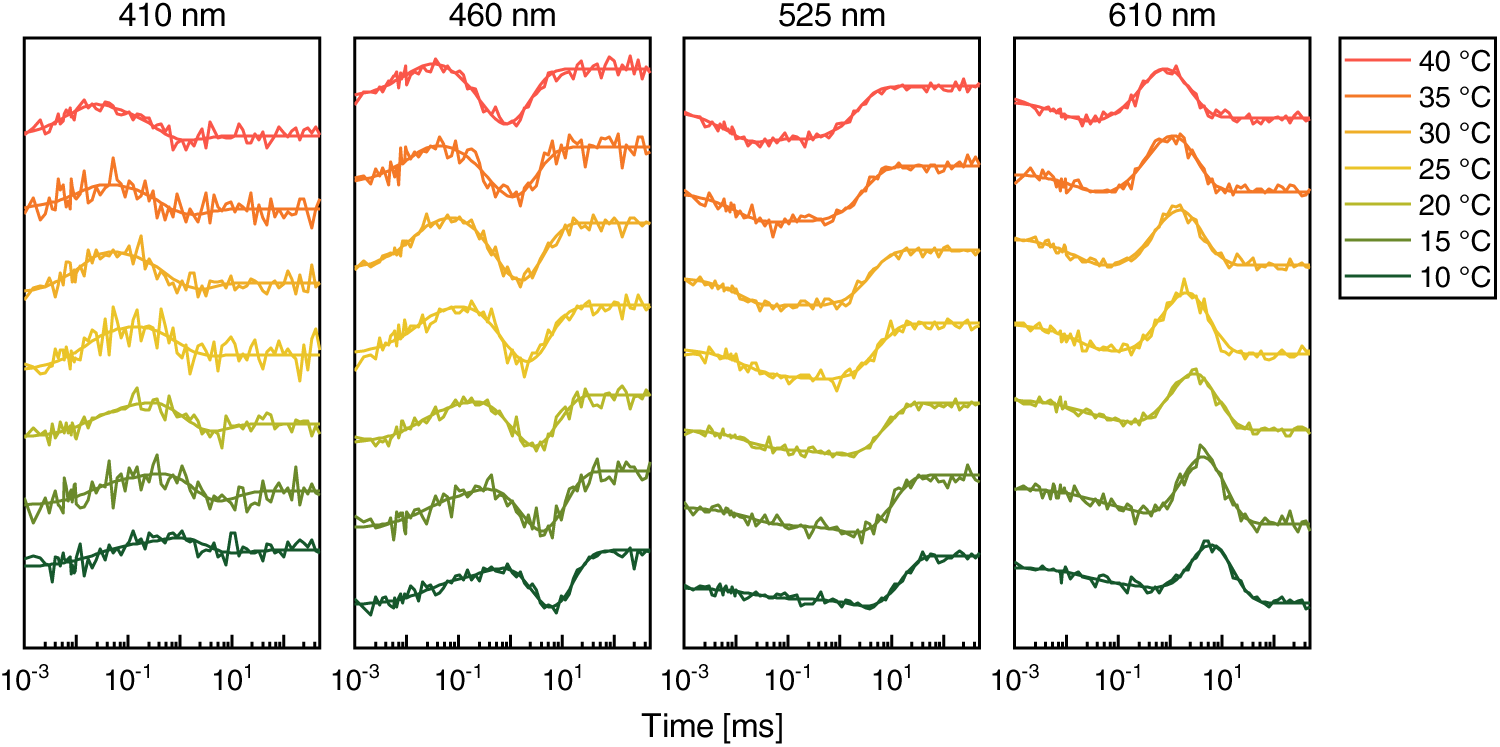
Flash photolysis measurement of KR2 WT at different wavelengths and temperatures.

The overall photocycle duration decreases with increasing temperature, from 10 to 40 °C by a factor of approximately six. To derive the number of kinetic components necessary for the target analysis we calculated lifetime density maps (LDMs) of all datasets (Figure S2). The LDMs revealed four main distributions, with the fastest one exhibiting the smallest change with temperature. The other three distributions move to earlier lifetimes with increasing temperature. This can be observed nicely by comparing the LDMs for 10 and 40 °C. Around 40 °C the second distribution becomes so fast, that it practically merges with the first distribution. The four distributions found in the LDMs were used as a starting point for the kinetic modeling.

### Target analysis

We performed global target analysis of all experimental datasets, i.e. each kinetic model was optimized on all datasets simultaneously. The kinetic rates at different temperatures were linked based on the transition state theory (see Materials and Methods). Different models were explored for their compatibility with the temperature dependent flash photolysis data. These included various branched reaction schemes and schemes with back rates. Due to the lack of excitation wavelength dependence of the flash photolysis data, all tested models were homogenous, and thus starting with 100% population of the first state. We also tried to introduce back rates, but this was challenging as it as the resulting energy barriers for the back reactions were unreasonable. Therefore, considering the transition state theory, the simple sequential model S_1_ → S_2_ → S_3_ → S_4_ → GS offered the most straightforward description of the temperature dependent experimental data.

The Eyring plot (Figure 3) shows that the second rate k_S2S3_, exhibits the highest temperature dependence, i.e. a 40-fold increase with rising temperature from 10 to 40 °C. With a decrease of ∼ three times, the first rate, k_S1S2_ exhibits the lowest temperature dependence. The rates of the remaining two transitions, k_S3S4_ and k_S4GS_ decrease eight and six folds, with rising temperature from 10 to 40 °C, correspondingly. The Eyring parameters for the different transitions are summarized in Table 1.

**Table 1:** Entropy, enthalpy and free energy values for the various photointermediate transitions in the KR2 photocycle.

| | $\Delta H$ [kJ/mol] | $T\Delta S$ [kJ/mol] | $\Delta G$ [kJ/mol] | driving force |
| --- | --- | --- | --- | --- |
| $S_1 \rightarrow S_2$ | 22.4 | -40.4 | 62.8 | $T\Delta S$ |
| $S_2 \rightarrow S_3$ | 89.2 | 22.3 | 66.9 | $\Delta H$ |
| $S_3 \rightarrow S_4$ | 47.2 | -25.8 | 73.0 | $\Delta H$ |
| $S_4 \rightarrow GS$ | 41.0 | -36.2 | 77.8 | $T\Delta S / \Delta H$ |

**Figure 3:**
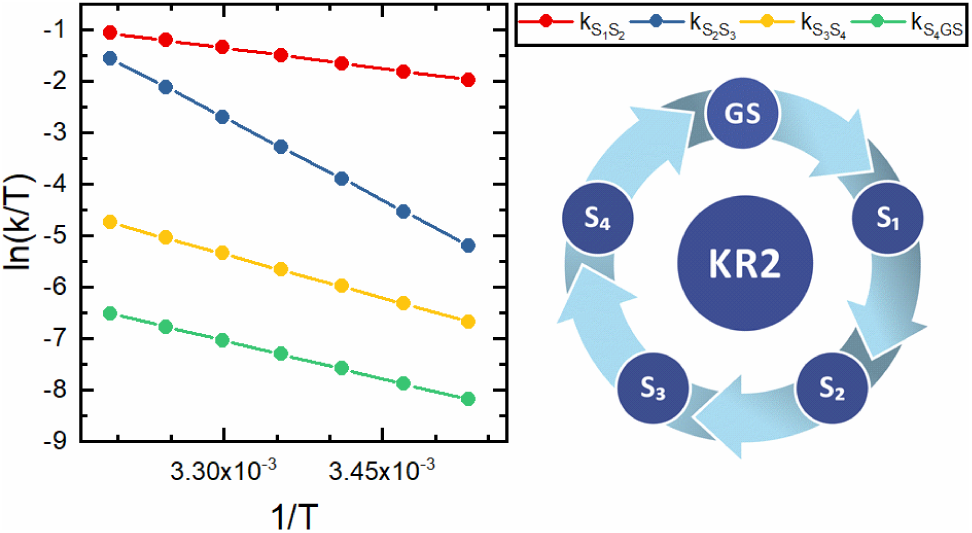
Eyring plot showing the temperature dependence of the single rates of the kinetic model.

While the S_1_-S_2_ transition appears to be driven by entropy, the S_2_-S_3_ and S_3_-S_4_ transitions are predominantly enthalpy driven. For the last step, the return to the ground state (S_4_-GS), entropy and enthalpy contribute equally to the driving force for the reaction.

The most prominent change in the transient populations (Figure 4) is observed for the S_2_ state. Its population is drastically reduced with higher temperature, while the S_3_ state reaches higher transient population. This picture follows the temperature dependence of the rates: while the S_1_ to S_2_ transition rate is only mildly increased at higher temperatures, the S_2_ to S_3_ transition rate is increased significantly. Therefore, at higher temperatures the S_2_ state is formed on a similar timescale, while it converts to the S_3_ state much faster, which effectively leads to a decrease of the overall S_2_ population at high temperatures.

**Figure 4:**
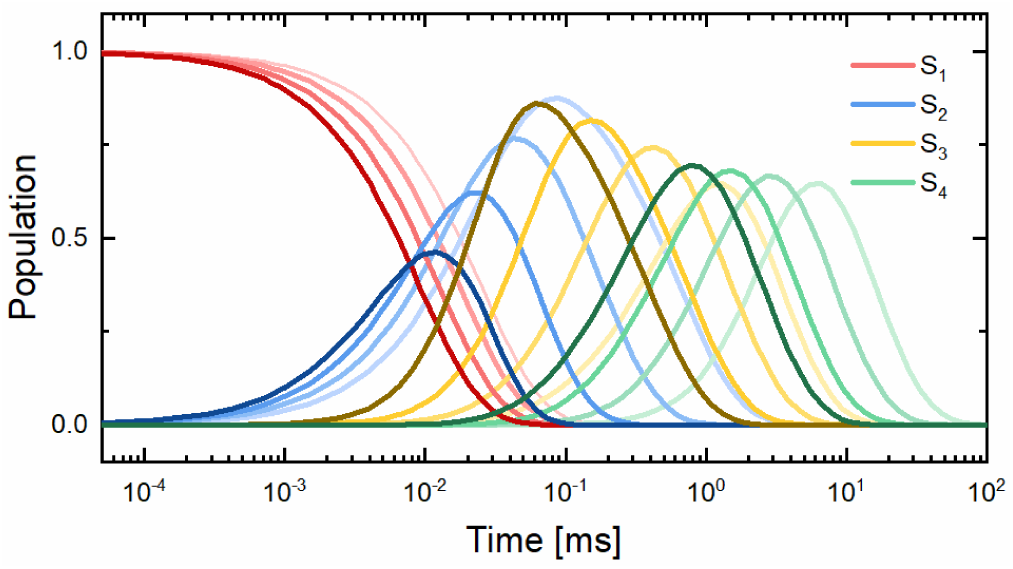
Transient population of the photocycle intermediates in the KR2 photocycle resulting from target analysis. Lighter colors represent lower temperatures; darker colors represent higher temperatures (10-40 °C in steps of 10 °C)

### Spectral correction

The kinetic modeling also yields the spectra of the involved states and their change at different temperatures. Since those spectra are obtained based on the absorbance difference data from flash photolysis experiments, they contain the GSB signal. Therefore, to determine the exact spectral position and band shape of the states, the GSB contribution needs to be subtracted.

The GSB signal can be derived from the ground state absorbance spectrum of KR2 (Figure S3). Therefore, only the retinal peak (λ_max_ = 525 nm) needs to be extracted and added to the difference spectra from the kinetic modeling. For this, it is crucial to take only the retinal peak contribution and not the whole absorbance spectrum (i.e. exclude contributions outside our spectral detection window originating from tryptophans and tyrosines around 280 nm). The peak at roughly 380 nm can contain contributions from the deprotonated retinal Schiff base and/or free retinal not bound inside the protein. For other retinal proteins like PR, the S_0_ – S_2_ transition has also been observed in this region^27^. However, for KR2 the extinction due to higher electronic transitions (S0-S2 or S0-S3) is very weak. Therefore, the contribution at 380 nm was not considered for the subtraction of the GSB.

Due to the size of the liposomes, the KR2 spectra in liposomes suffer from strong stray light, which makes extraction of the retinal peak more difficult than in, for instance, detergent samples. First, the absorbance spectrum must be corrected for the light scattering from the liposomes. Since the scattering is unique to every sample, it cannot be measured separately. To circumvent this problem, we used as a reference a KR2 spectrum from detergent solubilized sample, which does not suffer from light scattering and subtracted it from the spectrum of the liposome sample. This gave us access to the light scattering curve, which we fitted with a polynomial function. Subsequently, the fitted scattering curve was used to correct the liposome spectrum for the light scattering, resulting in an unperturbed steady state absorbance spectrum. From this corrected absorbance spectrum, we fitted only the retinal peak using a double Gaussian function. This gave us access to the unperturbed ground state bleach signal. This ground state bleach signal was finally subtracted from the difference spectra of the states in the kinetic modeling (Figure 5).

**Figure 5:**
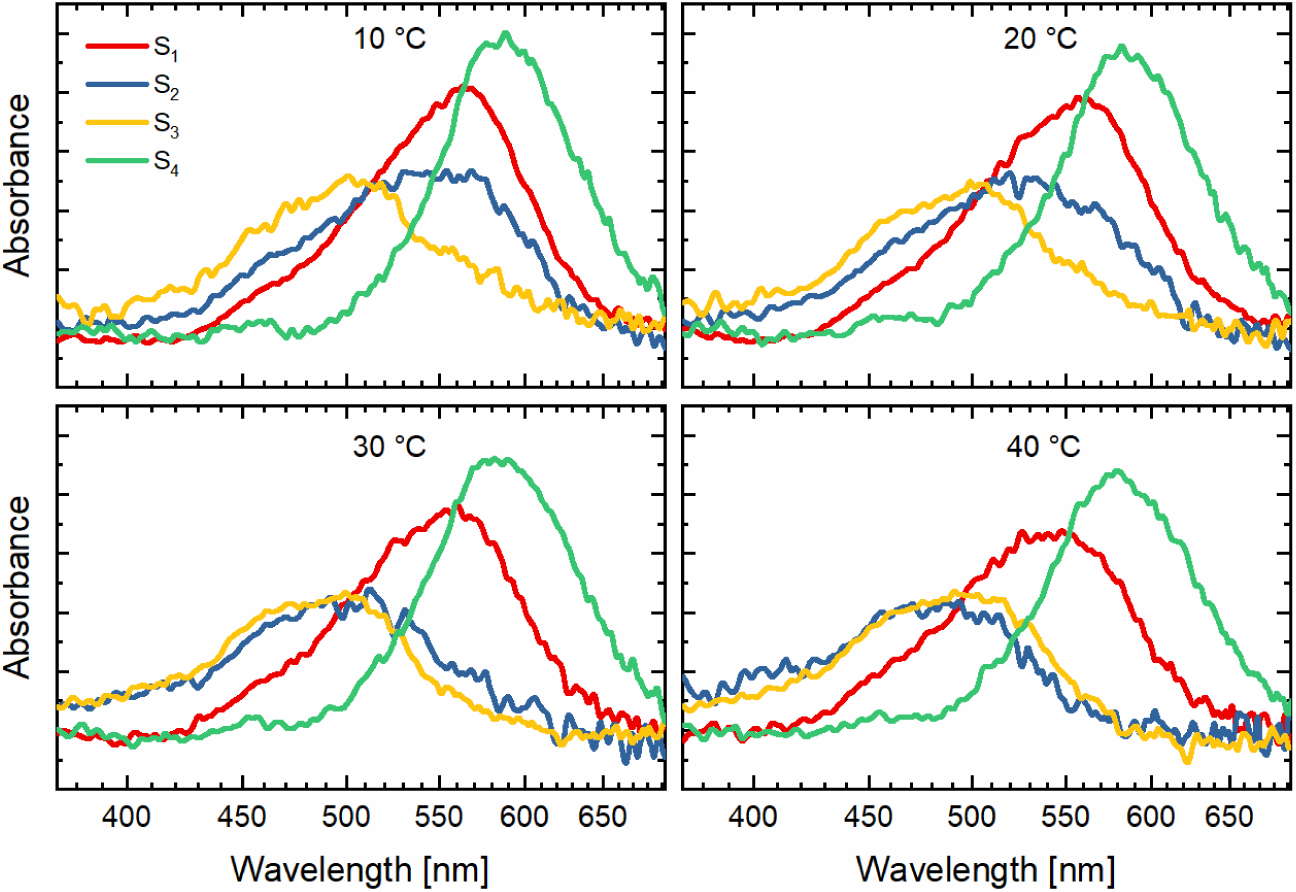
Evolution associated spectra of the various photocycle states at different temperatures.

The corrected spectra indicate a strong temperature dependence for the S_2_ spectrum, while S_1_, S_3_ and S_4_ spectra are not affected significantly. The S_2_ spectrum shifts from a more S_1_-like to a spectrum that is nearly identical to that of S_3_. At room temperature (20 °C), the S_2_ spectrum is positioned between the spectra of S_1_ and S_3_. The spectra of all states exhibit noticeable deviation from a pure Gaussian shape, therefore the question arises whether the transient states recovered from the kinetic modeling represent pure intermediates or their mixtures.

### Assignment of the kinetic components

To gain further insight into the kinetics of the KR2 photocycle we performed a spectral decomposition of the four kinetic states at each temperature.

Figure 6 shows the results from the spectral deconvolution at 20 °C. The first spectrum S_1_ actually consists of two Gaussian functions, which spectrally correspond to the K and the L intermediates. The second species S_2_ is represented by three Gaussian functions representing the K, L and M intermediates and S_3_ consists of the two Gaussian functions representing the L and M intermediates. Apart from the last species, which is always represented only by O, the intermediate with the furthest red shift, the composition of the other species slightly varies with temperature. The biggest changes can be seen for the second and third species, ranging from a mixture of K, L and M at 10 °C to a mixture of just L and M at 40 °C (Figure S4-S6). Finally, the fitted central wavelengths of the photointermediates are 563, 504, 449 and 584 nm for K, L, M and O, respectively. The spectrum of the M intermediate is the broadest with a FWHM of 48 nm while K is the narrowest with only 35 nm. The L and O intermediates have widths of 38 and 43 nm, correspondingly. Strikingly, the M intermediate, which is usually expected around 400 nm in other microbial rhodopsins,^16,28^ shows a comparably small blue shift in KR2. Figure 7 shows the spectra of the KR2 photointermediates resulting from the decomposition of the temperature dependent spectra obtained from the kinetic modeling.

**Figure 6:**
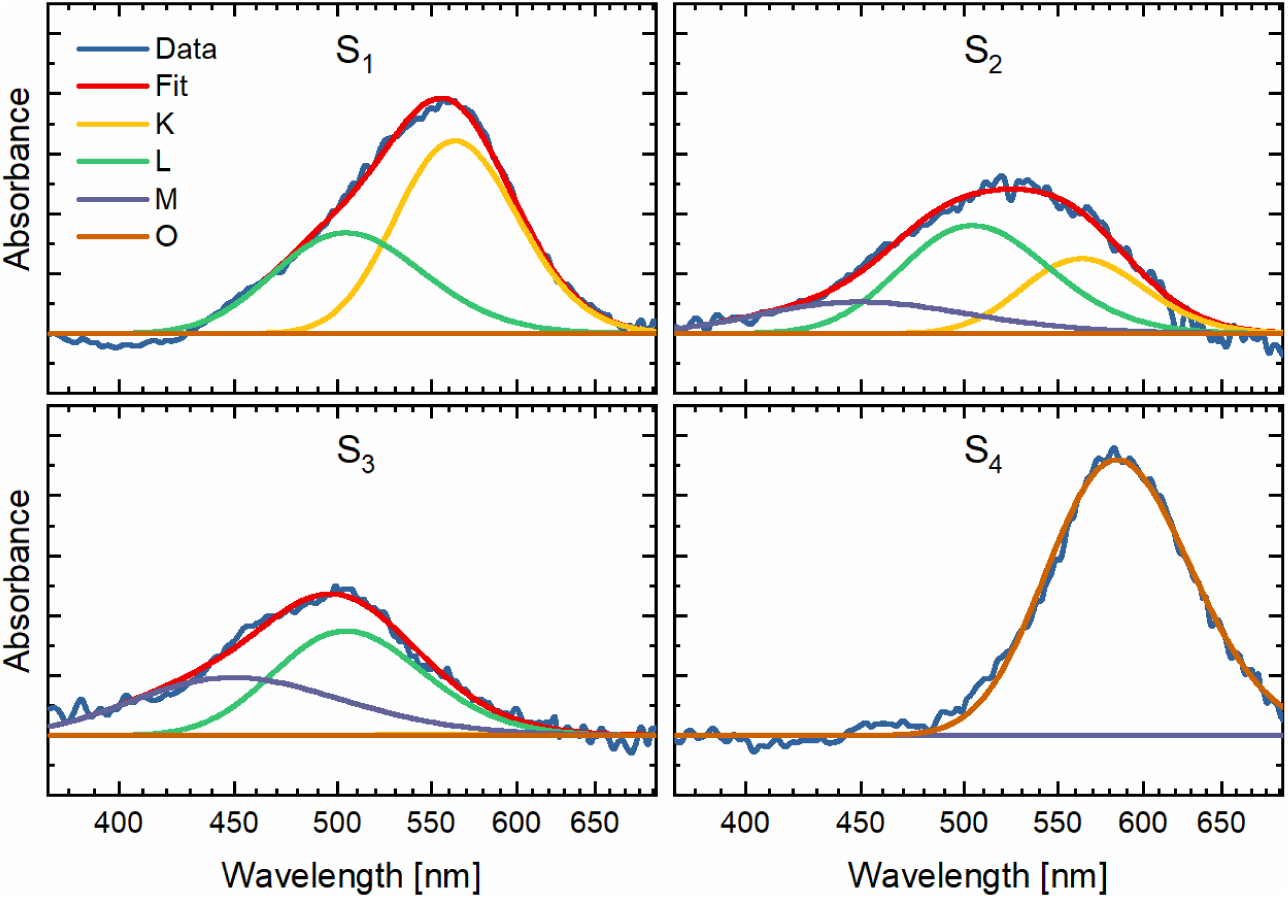
Spectral deconvolution of the four states at 20°C. The four panels show the data as well as the fit of the first, second, third and fourth species respectively. The spectral position as well as the width of the Gaussian functions for the deconvolution vary only little for the different temperatures.

**Figure 7:**
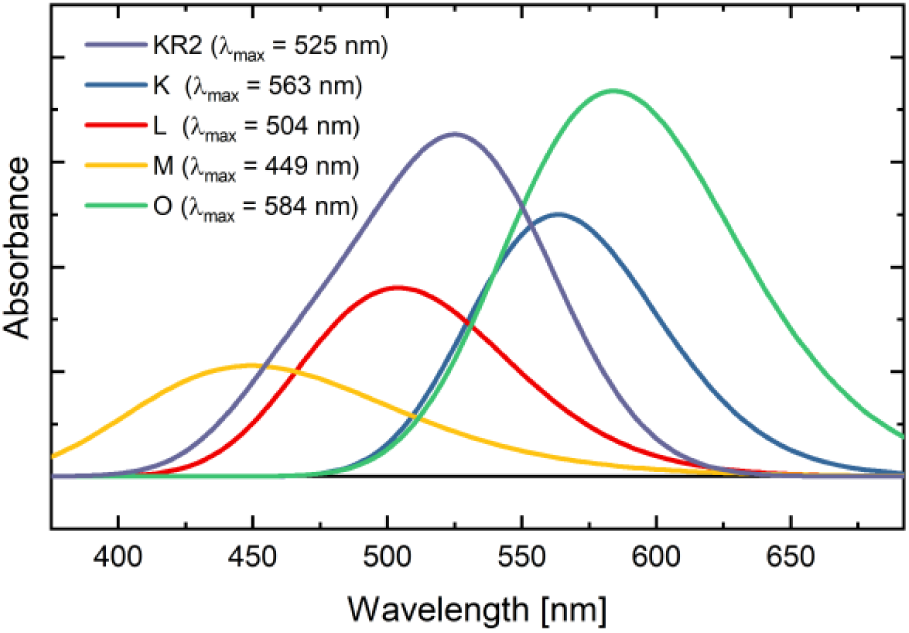
Spectra of the KR2 photointermediates and ground state derived from the decomposition of the temperature dependent spectra.

The necessity to use more than one Gaussian function for a satisfactory description of a given state in the kinetic modeling indicates that these states consist of equilibria between different photointermediates. Here, resolving these equilibria with the help of the target analysis turned challenging; rather only the transitions between these states were determined despite our efforts to utilize the transition state theory in the analysis. Therefore, we conclude that these equilibria between the photointermediates (contributing to the kinetic states in the model) are established very quickly. Our difficulties in resolving the kinetics in detail may also stem from differences in the reaction timescale of what is considered a “spectral” and a “protein” intermediate as the retinal and the protein evolve at a different pace. Nevertheless, the spectral decomposition allowed us to gain a more detailed picture of the temperature dependent kinetics.

Figure 8 shows the intermediate composition of the KR2 photocycle at different temperatures. At low temperatures, the photocycle starts in a K/L equilibrium state, going through two K/L/M equilibrium states of various compositions, then to the O intermediate, and finally back to the ground state. At 40 °C, the photocycle already contains a small amount of M intermediate in its initial state around 1 µs. After this initial state, the K intermediate decays and two equilibrium states with very similar spectra and slightly varying populations of the L and M intermediates (Figure S6) follow. Since their rates are also very close, they can also be fitted as one single state (Figure S7). Also, here, like in all cases, the pure O state is the last state before returning to the ground state. Since the O state always appears only by itself, not participating in any equilibria it is possible that the conversion to the O state is an irreversible step. This result is in agreement with a previously proposed mechanism, which suggests that the isomerization of the retinal chromophore back to the all-*trans* configuration and the reprotonation of the retinal Schiff base from D116 prevent the back flow of Na^+^ to the cytoplasmic side of the protein during O intermediate formation.^29^

**Figure 8:**
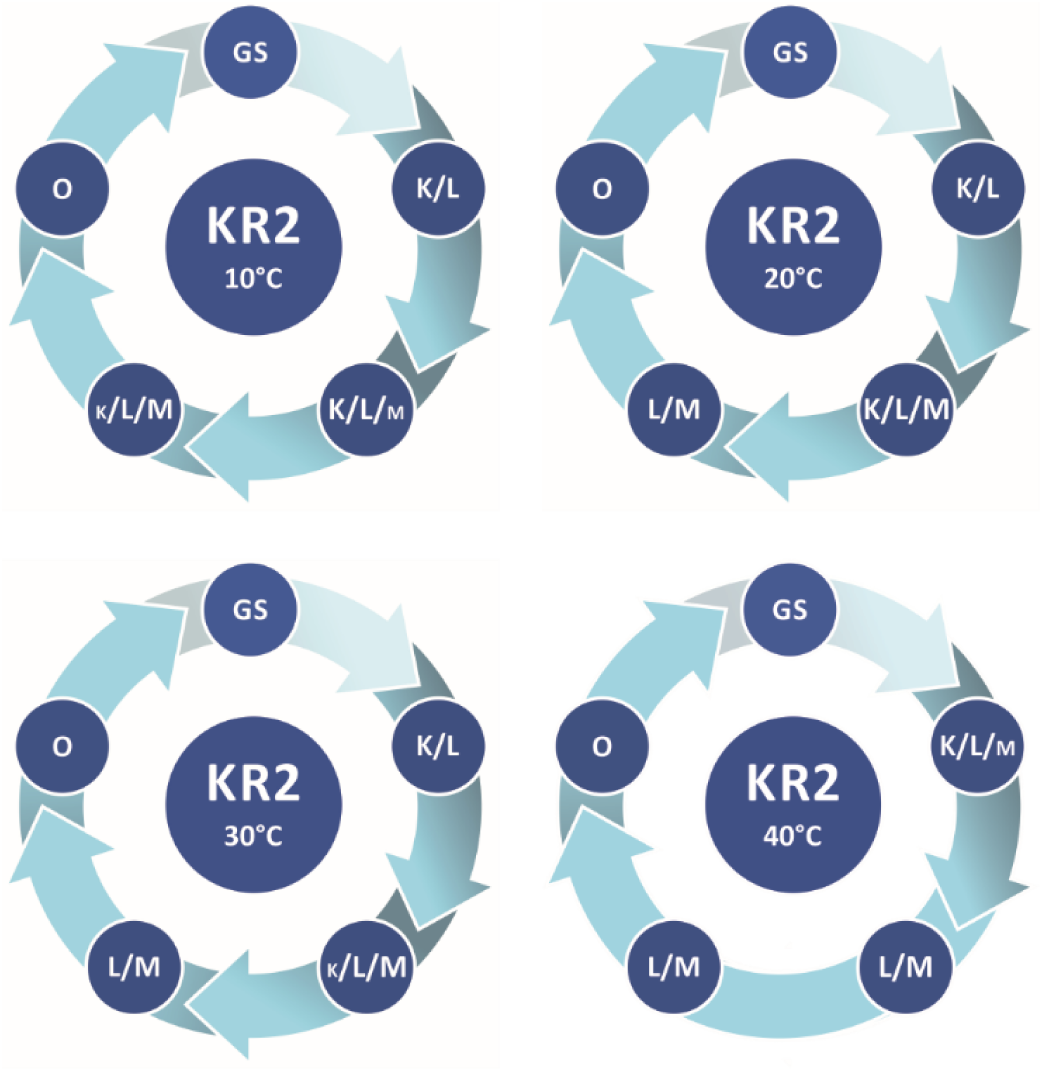
KR2 photocycle with equilibria of photointermediates at various temperatures.

## CONCLUSION

In this study, we investigated the temperature dependence of the KR2 photocycle. We could show that the different steps in the photocycle are affected to a different extent by the change in temperature. We observe several transient states that represent equilibria between the different photocycle intermediates. This result is in agreement with literature, where some of these equilibria could also be observed before.^12,22^ Our result shows that the transition from the L to the M intermediate is the step most affected by the temperature. While the overall photocycle is accelerated by a factor of six, the L-M transition is increased by a factor of 40. Using target analysis in combination with spectral decomposition the pure spectra of all the present photointermediates could be obtained, with their central wavelengths being 563 nm (K), 504 nm (L), 449 nm (M) and 584 nm (O). Gaining access to the dynamics, energies and spectra plays an important role in understanding the molecular mechanism of the protein and its use in optogenetics.

## ACKNOWLEDGEMENTS

The authors are supported by the Collaborative Research Center (SFB) 807 of the German Research Foundation (DFG).

## AUTHOR CONTRIBUTIONS

P.E., C.S., M.B. and J.W. conceived and designed the experiments. P.E. performed the experiments. P.E., C.S., M.B. and J.W. analyzed the data. J.S. and C.B. supplied the samples. P.E., C.S., M.B. and J.W. wrote the paper. All authors proofread and revised the manuscript.

